# Phosphoserine clusters as metal ion sensors for protein phase separation

**DOI:** 10.64898/2026.05.26.728006

**Authors:** Emilia T. Zakrzewska, Ahmed H. Mousa, Nicole Maurici, Dobroslawa Lewicka, Wiktor Kozminski, Alaji Bah, Rafal Augustyniak

## Abstract

Phosphorylation is a major regulator of biomolecular condensation, yet it remains unclear whether clustered phosphoserines can directly tune phase behavior via metal-ion coordination. Here, using solution NMR spectroscopy and human heterochromatin protein 1α (HP1α) as a model system, we show that stepwise phosphorylation of its N-terminal serine cluster generates a dynamic metal-responsive module that engages Mg²⁺, Ca²⁺, and Mn²⁺, whereas the unmodified protein shows little or no response. Metal coordination lowers the saturation concentration of phosphorylated HP1α, reshapes the temperature-dependent stability of its condensates, and modulates the effects of peptide regulators in an ion-specific manner. Our data support a model in which weak, transient metal-mediated contacts enhance intermolecular connectivity between phosphorylated HP1α molecules, promoting reversible condensation alongside canonical electrostatic interactions. These findings establish clustered phosphoserines as sequence-encoded metal-responsive elements that couple post-translational modification to the material properties of biomolecular condensates.

## Introduction

Condensate formation has emerged as a fundamental mechanism for organizing cellular biochemistry, enabling the formation of membraneless compartments that dynamically regulate biomolecular function without a surrounding lipid bilayer (1, 2). These condensates play essential roles in processes such as transcription, signaling, and genome organization, and their dysregulation has been linked to a range of diseases (3–8). Although the physical principles underlying phase separation are increasingly well understood, the molecular features that determine how protein condensation is regulated remain an active area of research (9, 10).

A key determinant of phase separation is intrinsic disorder. Intrinsically disordered regions (IDRs) of proteins enable transient, multivalent contacts that drive condensate formation, often through low-complexity sequences enriched in specific amino acids (2, 3, 11, 12). These contacts can be further tuned by post-translational modifications (PTMs), which alter interaction strengths and introduce new binding modes (7, 13, 14). Among these, phosphorylation is particularly widespread (15, 16). Rather than acting only at isolated sites, phosphorylation can also occur in clusters, generating regions of high local negative charge (17–19). In some cases, such clusters arise through hierarchical mechanisms in which initial phosphorylation events influence subsequent modifications at neighboring residues (17, 19, 20).

Despite their prevalence, the functional role of phosphoserine clusters remains incompletely understood. They are known to influence interactions with proteins and nucleic acids, particularly in nuclear and chromatin-associated proteins (19), yet their physicochemical contributions are less clear. The high density of negative charge suggests that these regions may engage in interactions beyond simple electrostatics. One possibility is that clustered phosphorylation creates multivalent interaction modules capable of attracting not only protein and nucleic acid partners but also metal ions through coordination. In some systems, such as caseins, clusters of phosphoserines are known to bind calcium ions and promote protein aggregation, highlighting their capacity to efficiently mediate intermolecular association (21, 22).

In parallel, the physicochemical environment has emerged as an important regulator of protein condensation. While monovalent salts primarily affect phase behavior through electrostatic screening (2), divalent metal ions can participate in more specific interactions, including direct coordination with biomolecules. A growing body of work shows that metal ions can modulate condensate properties (23). Calcium, for example, regulates phase separation in signaling complexes such as CaMKII, linking condensate formation to neuronal activity (24). Other ions, including copper and zinc, influence the phase behavior of proteins such as α-synuclein and tau, often shifting systems toward more condensed or aggregation-prone states (25, 26). Notably, engineered systems have demonstrated that even simple coordination motifs, such as hexahistidine tags, are sufficient to induce phase separation through metal-mediated clustering (27). These findings suggest that metal ions can act as active regulators of self-association rather than passive modifiers of ionic strength. However, the molecular basis of such effects remains poorly defined, particularly when post-translationally modified disordered regions are involved. It is therefore unclear which sequence features enable metal binding and how such interactions contribute to phase separation.

To address this question, we focused on heterochromatin protein 1α (HP1α), a conserved chromatin protein central to gene silencing and heterochromatin formation (28, 29). HP1α consists of a chromodomain (CD) that recognizes methylated histone H3 lysine 9, a chromoshadow domain (CSD) responsible for dimerization and partner recruitment, and three intrinsically disordered regions: the N-terminal extension (NTE), the hinge, and the C-terminal extension (CTE) (Fig. 1). HP1α self-associates both *in vitro* and in cells, forming condensates that resemble heterochromatin domains (30–32). These assemblies are driven primarily by interactions between post-translationally modified disordered regions and the folded domains, which together shape binding specificity and regulatory inputs (30, 33).

**Figure 1.**
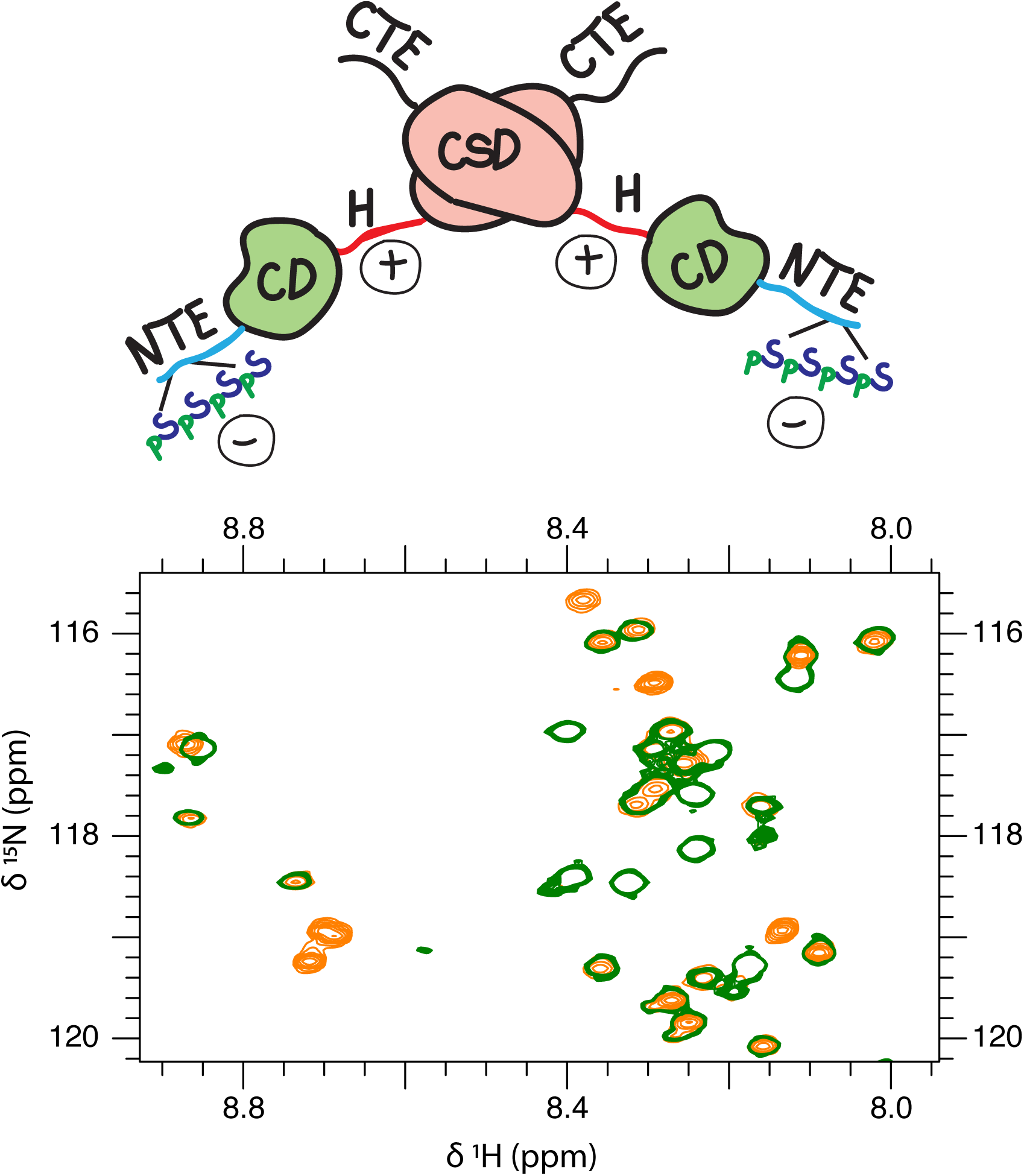
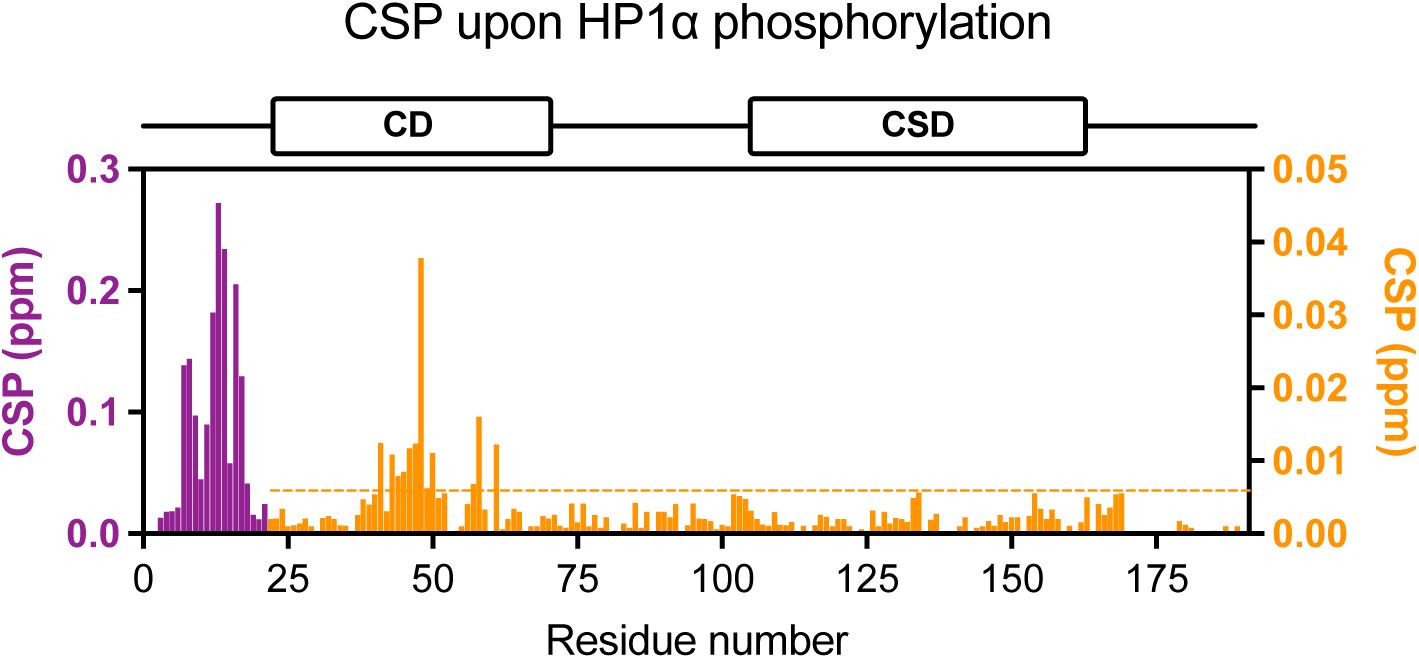
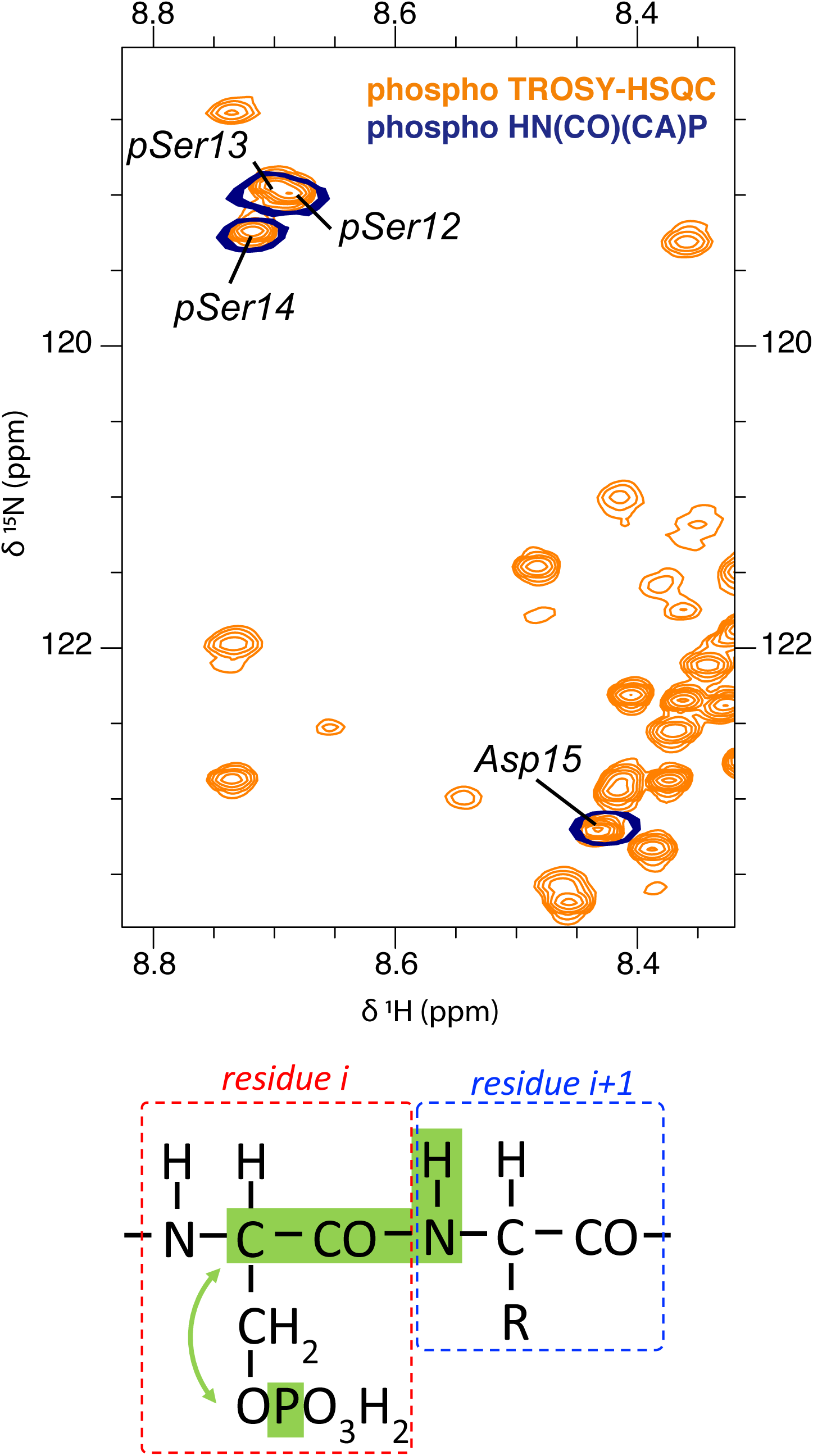
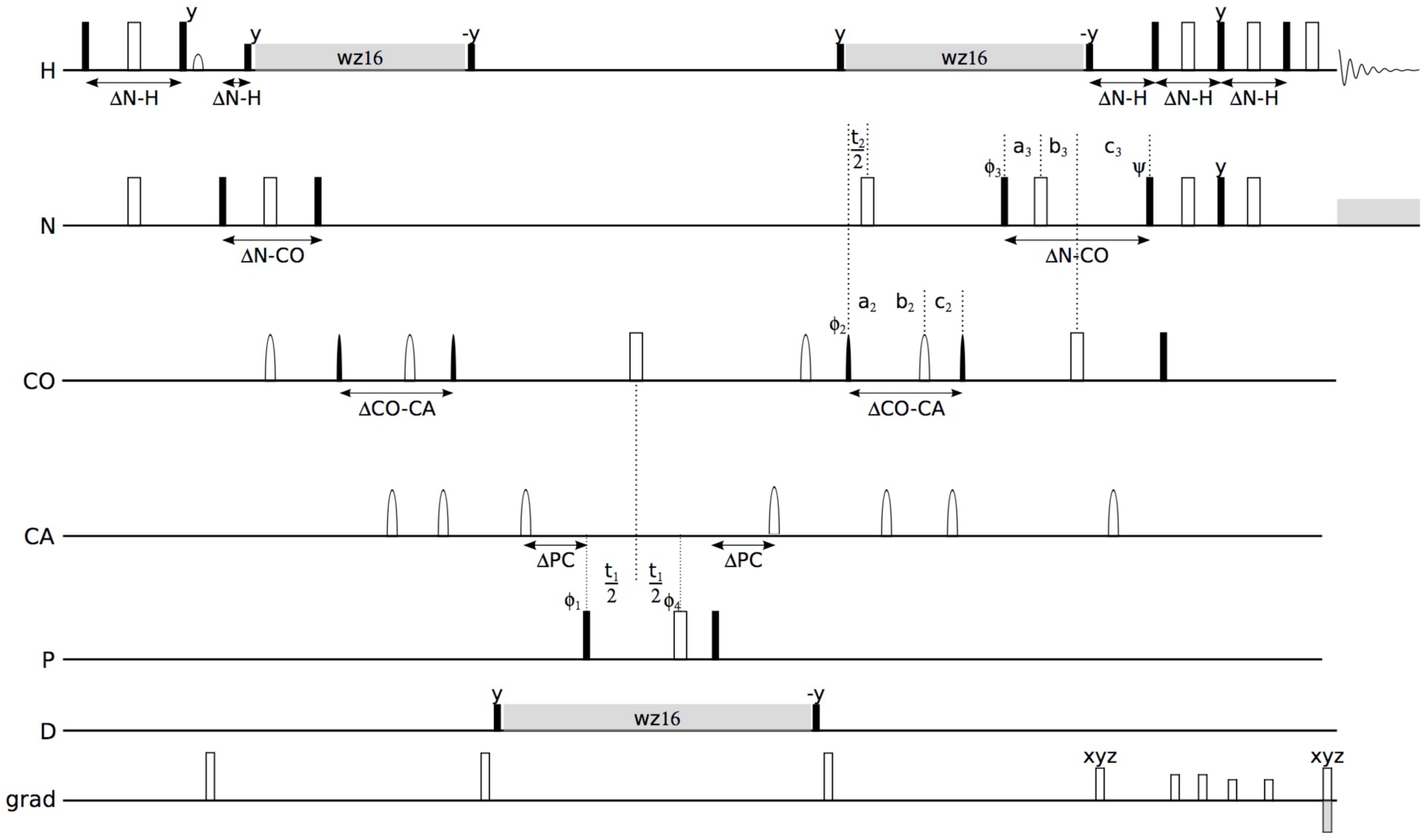
Identification of phosphorylation sites in HP1α and their structural consequences. (A,B) Comparison of NMR spectra of non-phosphorylated and phosphorylated HP1α showing phosphorylation-induced spectral changes within the N-terminal region. A schematic representation of HP1α domain organization and proposed intramolecular interactions is shown above the spectra. (C) Residue-specific chemical shift perturbations (CSPs) induced by phosphorylation mapped along the HP1α sequence, revealing strong local effects within the N-terminal extension and additional perturbations within the chromodomain region. (D,E) Identification of phosphorylated residues using HN(CO)(CA)P NMR experiments. Representative spectral regions and magnetization transfer pathways are shown. (F) Simplified pulse-sequence scheme used for phosphoserine detection. Together, the data demonstrate that phosphorylation of the N-terminal serine cluster induces both local and long-range structural effects in HP1α.

Importantly, phosphorylation of the N-terminal extension (NTE) is required for the phase separation, as the non-phosphorylated protein (non-phosHP1α) does not form droplets under comparable conditions (30, 31). This effect has been attributed to the introduction of negative charges, enabling interactions with positively charged regions, such as the hinge. At physiological pH, each phosphoserine contributes approximately two negative charges, whereas phosphomimetic substitutions such as glutamate provide only one. As a result, clusters of phosphoserines generate substantially stronger local charge than their mimetics, and indeed phosphomimetics fail to reproduce the phase behavior of phosphorylated HP1α (phosHP1α) (30). This suggests that phosphorylation does more than simply modulate electrostatics. The HP1α NTE contains a cluster of four consecutive serine residues (S11–S14) that are constitutively phosphorylated in cells (34). Whether this phosphorylated segment can support additional interactions, such as metal coordination, remains unclear.

Metal ions are also closely linked to higher-order chromatin organization. Mg²⁺ influences nucleosome interactions, chromatin compaction, and genome accessibility, and fluctuations in its concentration may affect nuclear phase behavior (35). Calcium can exert related but distinct effects, reflecting differences in coordination chemistry and binding specificity (36). Yet, despite the central role of HP1α in heterochromatin formation, the potential interplay between HP1α phosphorylation and metal ions has not been explored.

To investigate these processes at atomic resolution, we turned to nuclear magnetic resonance (NMR) spectroscopy. NMR chemical shifts are highly sensitive to the local environment and provide direct insight into structure, dynamics, and interactions at the residue level (37, 38). This is particularly valuable for intrinsically disordered proteins, where transient contacts and conformational heterogeneity are central to function (39, 40). In addition, NMR enables direct observation of post-translational modifications in intact proteins and in real time. For example, Selenko and co-workers demonstrated that phosphorylation can be monitored with residue-specific resolution *in vitro*, in cell extracts, and even in living cells, allowing closely spaced sites to be distinguished and their sequential modification resolved (41). Furthermore, NMR approaches developed to probe conformational exchange and low-populated states have been successfully applied to biomolecular condensates, providing residue-level insight into interactions within phase-separated systems (42, 43). These capabilities make NMR uniquely suited to connect sequence, dynamics, and chemical modification in complex systems.

Here, we asked whether clustered phosphorylation can directly couple to metal coordination to regulate phase separation. Using HP1α as a model system, we demonstrate that a phosphoserine-rich segment of the NTE functions as a dynamic metal-binding module that selectively engages divalent cations. This interaction is strictly dependent on phosphorylation, establishing a direct link between post-translational modification and metal ion recognition. We find that metal coordination reshapes the phase behavior of HP1α and rewires the effects of regulatory interactions. These results support a model in which multiple phosphorylated HP1α molecules transiently share metal ions, generating dynamic multivalent cross-links that drive phase separation independently of canonical electrostatic interactions. This work establishes phosphoserine clusters as chemically defined modules for metal-dependent regulation of biomolecular condensates.

## Results

### NMR reveals site-specific phosphorylation of HP1α

To characterize the effect of phosphorylation on HP1α, we compared ¹H–¹⁵N TROSY HSQC spectra of the unmodified protein with those of the protein after *in vitro* phosphorylation by casein kinase (CK2). The reaction was followed by mass spectrometry, which confirmed incorporation of four phosphate groups (Fig. S1). The sample was then dialyzed into the same buffer used for non-phosphorylated HP1α (50 mM HEPES, 150 mM KCl, 2 mM TCEP, pH 7.0), thereby removing ATP and MgCl₂ prior to NMR analysis. Under these matched solution conditions, phosphorylation induces extensive spectral changes dominated by the N-terminal extension (Fig. 1B-C). These chemical shift perturbations (CSPs) are not restricted to the S11–S14 phosphorylation site itself but extend across nearly the entire NTE, indicating that phosphorylation reorganizes the conformational ensemble of this region rather than acting as a purely local modification, consistent with previous work on a truncated version of HP1α (44). Much smaller but reproducible changes are also detected in the chromodomain, whereas the hinge and the remainder of the protein are essentially unaffected. Although the hinge contains a CK2 consensus motif in human HP1α (S97 followed by D99 and D100), neither the NMR data nor mass spectrometry provides evidence for modification outside the NTE.

Within the chromodomain, the largest CSPs are observed for residues Y20, V21, H48, N49, T50, E52, L57, D58, and E61 (Fig. 1C). In the structure of the HP1α chromodomain bound to an H3K9me3 peptide (PDB code 3FDT), these residues map directly onto the histone-binding surface (45). Y20, V21, E52, L57, and D58 make direct contacts with the peptide, while H48, N49, and T50 contribute to recognition of the histone tail through contacts with the trimethylated lysine K9. Thus, the residues perturbed by phosphorylation define the same surface that engages the H3K9me3 mark, indicating that the NTE and the CD are not independent in solution but communicate through long-range intramolecular contacts in which phosphorylation of the NTE alters the histone-binding interface of the CD.

To obtain residue-specific information on phosphorylation sites, we developed an NMR experiment that exploits the scalar coupling between the Cα nucleus and the phosphate group in phosphoserine (³J_Cα–P_ ≈ 5–7 Hz). Similar couplings have been used to detect phosphorylated residues using spectral editing approaches, such as ¹³C–³¹P spin-echo difference experiments (46). However, these methods rely on indirect detection and are difficult to extend to higher-dimensional experiments required for intrinsically disordered or hyperphosphorylated proteins, where severe spectral overlap limits the utility of simple 2D approaches.

In our implementation, magnetization is transferred from the amide proton through the backbone to Cα and then to the phosphate group before being returned for detection. Because the experiment is compatible with non-uniform sampling, it can be extended to higher-dimensional datasets—up to five correlated chemical shift dimensions—while still maintaining reasonable experimental times. When displayed in a ¹H–¹⁵N plane, the data reduce to an HSQC-like pattern containing only four signals present (Fig. 1B). These peaks correspond to residues S12, S13, S14, and E15, reflecting the *i–*1 transfer pathway illustrated in Fig. 1D, and reporting on phosphorylation of the preceding serine side chains. In addition, the experiment provides direct access to ³¹P chemical shifts. A ¹H–³¹P projection (Fig. S2) reveals four distinct phosphate resonances, consistent with four phosphoserine sites and indicating that each phosphate group experiences a slightly different chemical environment.

### Hierarchical phosphorylation of the HP1α N-terminal cluster

To monitor phosphorylation in real time, we recorded a series of ¹H–¹⁵N TROSY HSQC spectra after addition of CK2, ATP, and Mg²⁺. These data reveal a clear biphasic behavior (Fig. 1E). Residues flanking the serine cluster, including T8, A9, and D10, show disappearance of the initial signals accompanied by the appearance of new peaks, which are subsequently replaced by a second set of resonances as the reaction proceeds. This behavior is inconsistent with a simple concerted process and instead points to the formation of at least one intermediate state.

Direct insight into the first phosphorylation event is provided by the HNCoCaP experiment. The earliest signal to appear corresponds to E15, reporting on phosphorylation of S14. This resonance initially accumulates and then shifts to a different position as the reaction progresses, indicating that the local environment of pS14 changes as neighboring residues are further modified. Quantitative analysis of signal intensities (Fig. 1F) supports this interpretation. The original S14 signal disappears rapidly, and a single sharp pS14 resonance appears in its place. This early signal reaches a maximum after ∼4–6 hours and then begins to decay. At this stage, the phosphoserine region becomes populated by a series of weak peaks that likely reflect a mixture of partially phosphorylated intermediates. As the reaction proceeds, these transient signals give way to a smaller number of well-resolved pS resonances corresponding to the final state.

Mutational analysis further clarifies the mechanism. Substitution of S14 by alanine abolishes phosphorylation, whereas S11A and S12A/S13A mutants retain the biphasic behavior (Fig. S3). These results show that phosphorylation of the HP1α N-terminal serine cluster proceeds through a hierarchical mechanism initiated by a rapid priming event at S14, followed by a slower phase in which S11, S12, and S13 are modified in the context of an already S14-phosphorylated protein. While the precise order of these latter events cannot be resolved from the present data, the persistence of intermediate states and redistribution of pS signals indicate that multiple partially phosphorylated species coexist during the reaction.

### Phosphoserine cluster mediates weak and dynamic metal ion interactions

We next examined whether the phosphorylated N-terminal region of HP1α can interact with divalent metal ions. This question was prompted by the observation that the ¹H–¹⁵N HSQC spectrum of phosphorylated HP1α recorded directly after the CK2 reaction differed from that obtained after buffer exchange, in which ATP and MgCl₂ were removed. This suggested that components of the phosphorylation mixture, most notably Mg²⁺, interact with the phosphorylated protein.

To test this directly, we performed NMR titrations of MgCl₂ into buffer-exchanged phosHP1α under high-salt conditions (200 mM KCl) and at 25 °C, where phase separation is suppressed, and monitored chemical shift perturbations (Fig. 2A). The resulting changes are highly localized. The largest CSPs are confined to the NTE encompassing the phosphoserine cluster (pS11–pS14) and adjacent residues. Modest perturbations are observed for H48 and nearby residues in the chromodomain. Histidine side chains are known to coordinate divalent metal ions, and this localized response likely reflects a weaker, secondary interaction. No detectable changes occur elsewhere in the protein (Fig. 2B). By comparison, the non-phosphorylated protein does not exhibit comparable perturbations under the same conditions, indicating that the interaction is strictly dependent on phosphate groups. The titration curves are smooth and gradual, without clear saturation, consistent with weak interactions in fast exchange on the NMR timescale (Fig. 2C).

**Figure 2.**
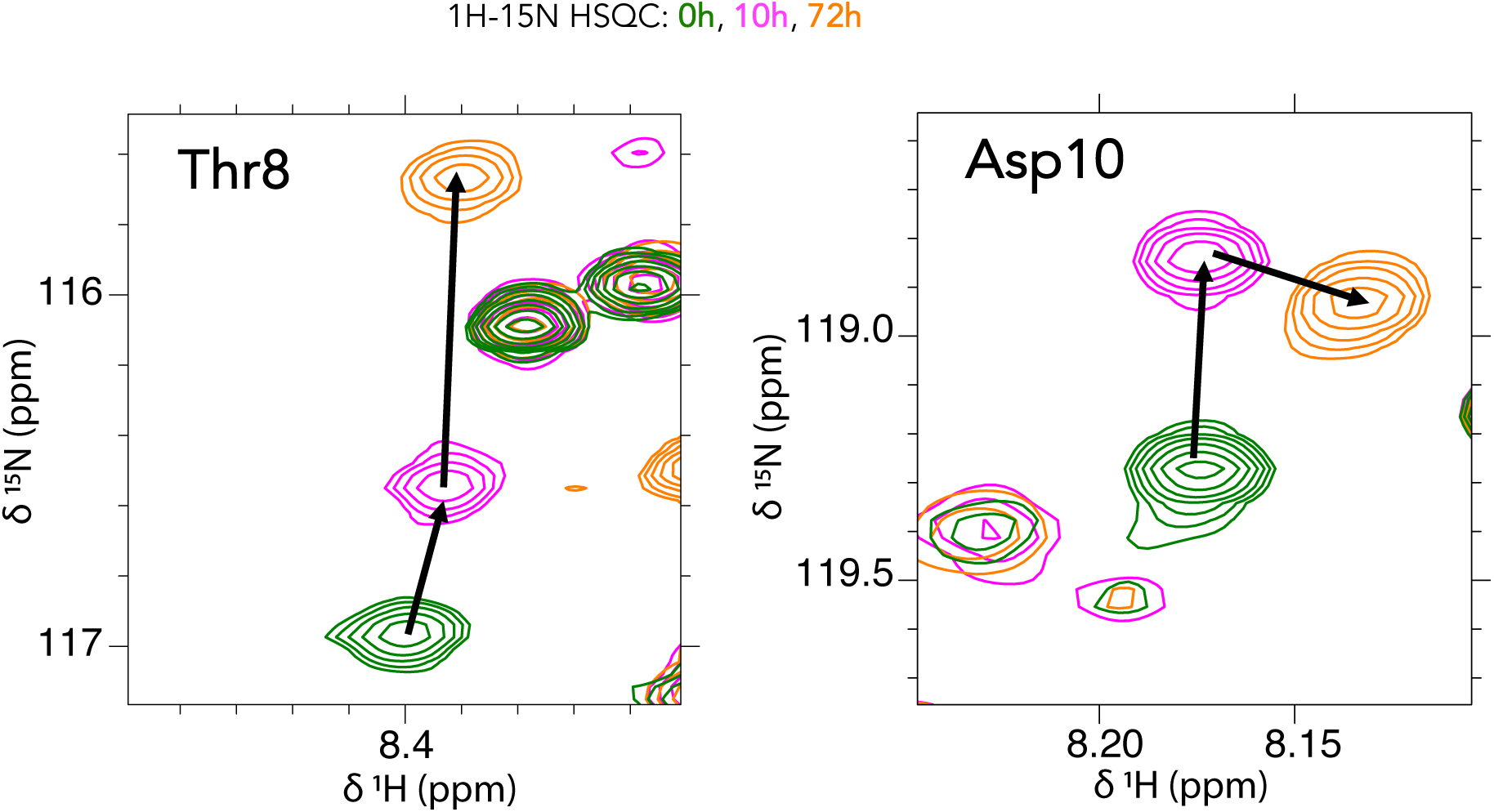
Time-resolved monitoring of HP1α phosphorylation by NMR spectroscopy. Sequential ¹H–¹⁵N HSQC spectra recorded during the CK2 phosphorylation reaction reveal the progressive appearance of phosphoserine resonances and disappearance of signals corresponding to the non-phosphorylated state. Expanded spectral regions illustrate the time-dependent evolution of selected residues within the N-terminal extension. The data demonstrate that phosphorylation of HP1α can be directly monitored in real time at residue-specific resolution.

Because the phosphorylated N-terminus constitutes a multivalent cluster rather than a single, well-defined binding site, the system cannot be described by a simple 1:1 binding model. We therefore analyzed the titration curves using an empirical saturating function and report apparent EC50 values rather than microscopic dissociation constants (see Supplementary Methods). Individual fits yielded similar concentration dependence but differed in amplitude, whereas a global fit with a shared EC50 parameter provided a consistent description across the phosphoserine cluster. pS11 showed a modest deviation from this behavior, consistent with its position at the edge of the phosphorylated segment. This analysis yielded an apparent EC50 of 8.4 ± 0.7 mM for Mg²⁺.

We then examined the interaction of phosHP1α with Ca²⁺. Titration with CaCl₂ produces a similar pattern of localized CSPs, largely confined to the phosphoserine cluster and its immediate surroundings (Fig. 2C). Compared with Mg²⁺, the magnitude of the perturbations is substantially larger, and the response is more heterogeneous across individual phosphoserines. In particular, pS13 shows a markedly stronger response than the other sites, indicating non-uniform coordination across the cluster. No comparable perturbations are observed for H48, suggesting that Ca²⁺ interacts predominantly with the phosphorylated NTE rather than engaging additional sites in the chromodomain.

Global analysis of the titration data yields an apparent EC50 of 4.7 ± 0.2 mM for Ca²⁺, which is lower than that for Mg²⁺. While both ions interact with the phosphorylated N-terminus in a similar overall manner, their effects differ in both strength and residue-specific response, consistent with distinct modes of coordination within the multivalent phosphoserine cluster.

To further probe the nature of the interaction between divalent cations and the phosphorylated N-terminal region, we turned to paramagnetic relaxation enhancement (PRE) using Mn²⁺. Addition of only 1 µM MnCl₂ to 300 µM phosphorylated HP1α results in the disappearance of the phosphoserine signals, accompanied by strong attenuation of nearby backbone resonances within the N-terminal extension (Fig. 2D). In contrast, no comparable effects are observed for the non-phosphorylated protein, confirming that the interaction requires the presence of phosphate groups.

Given the strong distance dependence of PRE, this observation indicates that Mn²⁺ approaches the phosphoserine cluster at short distances in solution. The magnitude of the effect at such low sub-stoichiometric Mn²⁺ concentrations does not reflect tight binding but rather the high sensitivity of PRE to transient close encounters. Even a single short-lived approach during an NMR acquisition can strongly attenuate the signal, and repeated sampling of the phosphorylated region over multiple scans leads to cumulative signal loss. Thus, a small number of Mn²⁺ ions can effectively probe a large population of HP1α molecules without forming stable complexes. This behavior is consistent with a dynamic mode of coordination in which Mn²⁺ can transiently bridge different protein molecules over time.

Together with the smooth CSP titrations observed for Mg²⁺ and Ca²⁺, these data support a model in which divalent metal ions interact with the phosphoserine cluster through weak, rapidly exchanging coordination events. Rather than forming stable stoichiometric complexes, these transient interactions are well suited to promote reversible intermolecular contacts and provide a mechanism by which metal ions facilitate phase separation of HP1α.

### Divalent metal ions promote phase separation of phosphorylated HP1α

To assess whether divalent metal ions induce phase separation of phosHP1α, we first examined the protein behavior upon addition of MnCl₂ under low-salt conditions. At room temperature, a solution of 300 µM phosHP1α in LLPS buffer (50 mM HEPES, 75 mM KCl) was initially transparent but became visibly turbid upon addition of 5 mM MnCl₂ (Fig. S4). This effect was fully reversible upon addition of EDTA, indicating that droplet formation depends on metal coordination rather than irreversible aggregation. Microscopy confirmed the formation of liquid-like droplets (Fig. 3A). Similar behavior was observed for Mg²⁺ and Ca²⁺, although the extent of droplet formation depended on ion identity, concentration, and temperature.

**Figure 3.**
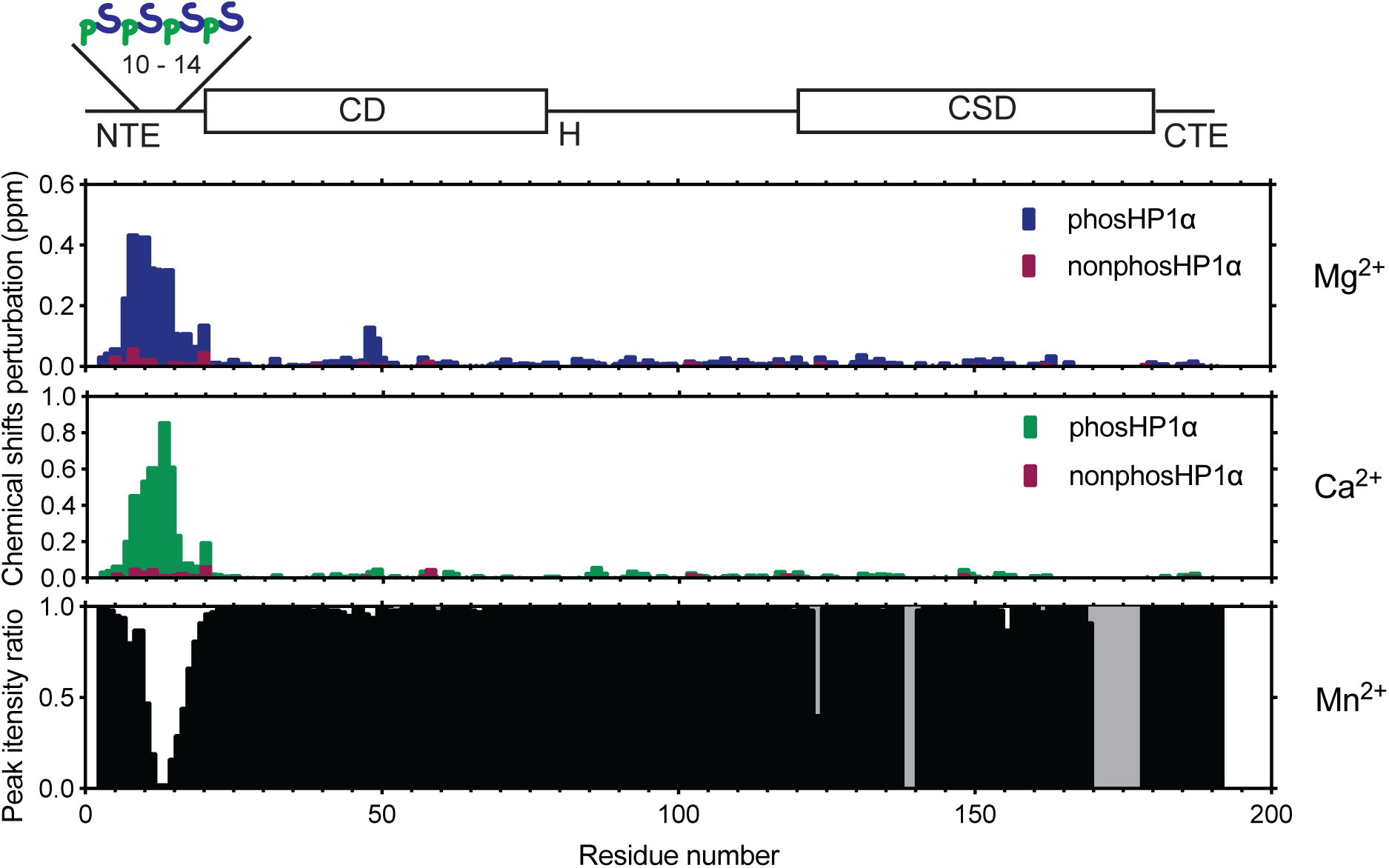
Phosphorylated HP1α directly interacts with divalent metal ions. Chemical shift perturbation and signal intensity analyses showing the effects of Mg²⁺, Ca²⁺, and Mn²⁺ on phosphorylated and non-phosphorylated HP1α. Mg²⁺ and Ca²⁺ induce localized perturbations primarily within the phosphorylated N-terminal serine cluster, whereas Mn²⁺ causes strong signal attenuation consistent with paramagnetic relaxation enhancement effects. In contrast, non-phosphorylated HP1α exhibits little or no response under comparable conditions. The data support direct and phosphorylation-dependent interactions between HP1α and divalent metal ions.

**Figure 4.**
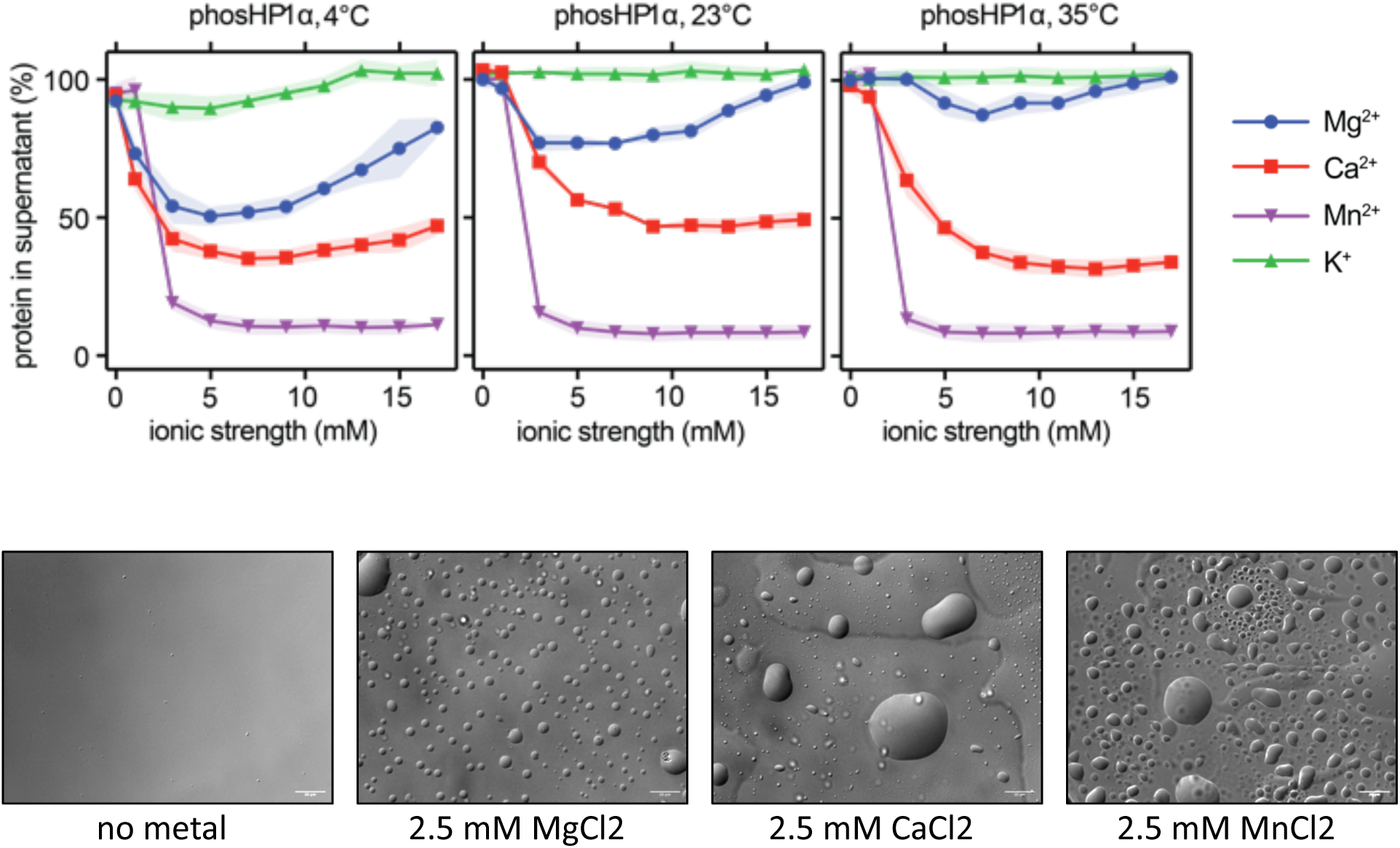
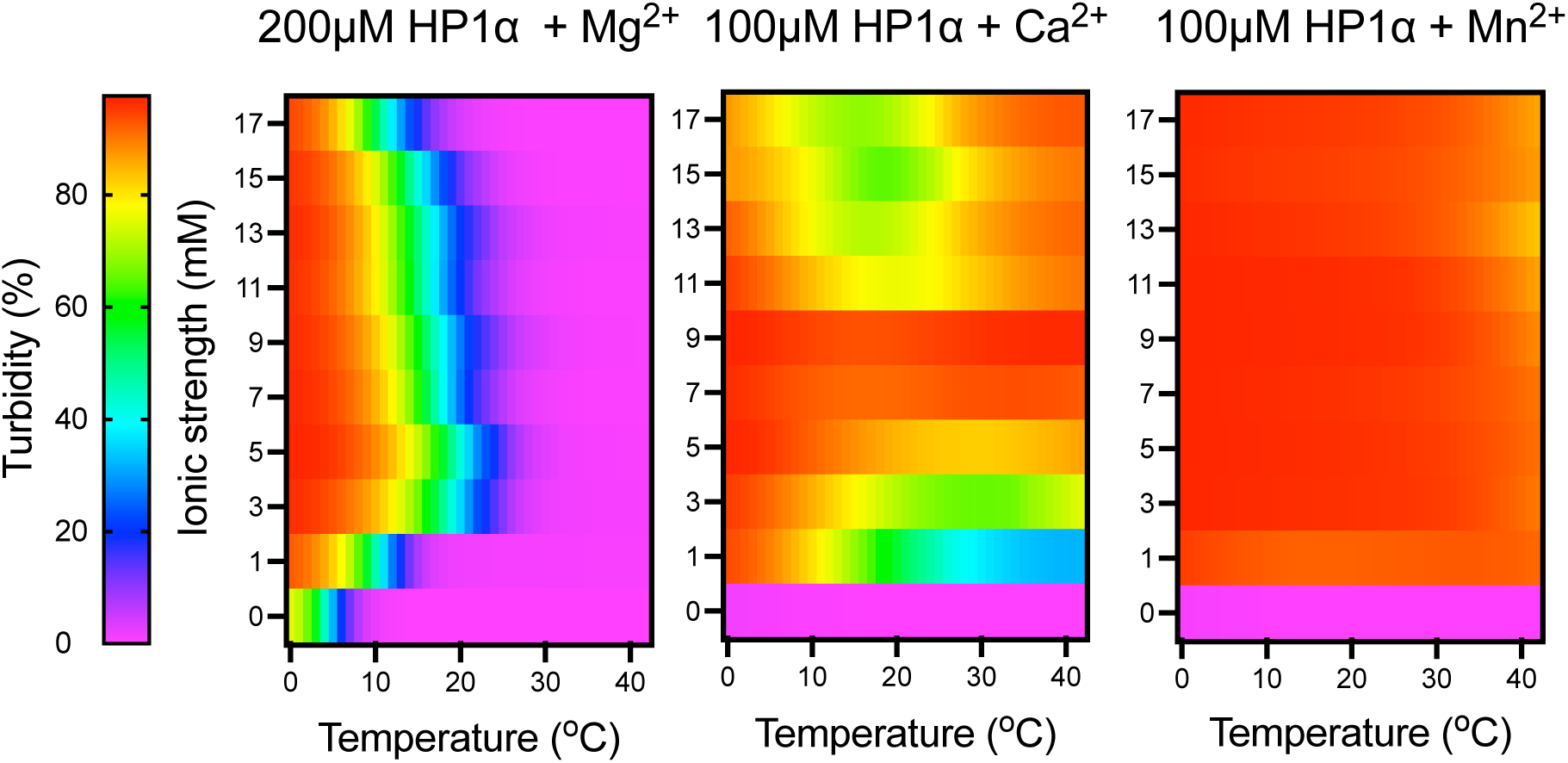
Metal ions promote phase separation of phosphorylated HP1α in an ion-specific manner. (A) Centrifugation-based depletion assays showing the effects of Mg²⁺, Ca²⁺, Mn²⁺, and monovalent salt controls on HP1α phase separation at different temperatures. (B) Representative microscopy images illustrating metal ion-induced droplet formation. (C) Temperature-dependent turbidity heatmaps showing the effects of increasing concentrations of Mg²⁺, Ca²⁺, and Mn²⁺ on the stability and phase behavior of HP1α condensates. The data reveal distinct ion-specific effects on condensate formation and thermal stability.

To quantify these effects, we performed spin-down assays to determine the fraction of protein remaining in the dilute phase. PhosHP1α was incubated with increasing concentrations of MgCl₂, CaCl₂, or MnCl₂ at defined temperatures, followed by centrifugation under the same conditions. Protein concentration in the supernatant was determined by absorbance at 280 nm and expressed relative to the no-metal control at each temperature.

To distinguish specific metal effects from nonspecific electrostatic screening, KCl was included as a monovalent reference. To enable direct comparison between mono– and divalent salts, the x-axis was expressed as the ionic strength contribution of the added salt (ΔI), such that 1 mM KCl contributes 1 mM ionic strength, whereas 1 mM MgCl₂, CaCl₂, or MnCl₂ contributes 3 mM.

As expected for electrostatic screening, increasing KCl concentration did not promote phase separation and instead increased the fraction of protein remaining in the supernatant. At 4 °C, phosHP1α showed only limited baseline depletion, which was progressively suppressed at higher ionic strength. No phase separation was detected at 23 °C or 35 °C.

In contrast, Mg²⁺ induced a pronounced, non-monotonic response. At 4 °C, increasing MgCl₂ concentration enhanced phase separation at intermediate concentrations, followed by partial reversal at higher concentrations. Similar trends were observed at 23 °C and 35 °C, although with reduced magnitude. Notably, Mg²⁺ enabled phase separation under conditions where the no-metal sample remained fully soluble.

Ca²⁺ produced a distinct response. At 4 °C, calcium induced stronger depletion from the supernatant than Mg²⁺, indicating more efficient partitioning into the dense phase. This effect was less susceptible to reversal at higher concentrations. Strikingly, the temperature dependence was non-monotonic: phase separation was reduced at 23 °C but increased again at 35 °C, yielding substantial depletion comparable to low-temperature conditions. This behavior is consistent with the coexistence of upper and lower critical solution temperature regimes (UCST and LCST) (47).

Mn²⁺ exhibited the most pronounced effect. Above a threshold concentration (∼3 mM), phase separation was strongly enhanced at all temperatures, with only a small fraction of protein remaining in the supernatant. Below this threshold, little effect was observed. This sharp transition is consistent with a collective mechanism in which Mn²⁺ promotes intermolecular connectivity and stabilizes the dense phase.

Overall, these results demonstrate that divalent metal ions strongly modulate the phase behavior of phosphorylated HP1α in a manner that depends on ion identity, concentration, and temperature. In combination with NMR data indicating weak and dynamic interactions with the phosphoserine cluster, these findings support a model in which metal ions act as transient crosslinkers that enhance intermolecular interactions and drive phase separation.

### Metal ions reshape the temperature-dependent phase behavior of HP1α

To further characterize how metal ions influence phosHP1α phase behavior, we monitored turbidity during controlled temperature ramps. Unlike spin-down assays, which probe phase separation under fixed conditions, this approach captures how the stability of the condensed phase varies with temperature. PhosHP1α samples were subjected to a gradual temperature increase, and turbidity was recorded as a function of temperature. Metal ions were added at concentrations matching those used in spin-down assays, allowing direct comparison. The resulting data are presented as heatmaps, where color intensity reflects the extent of phase separation across temperature and metal ion concentration.

At a protein concentration of 200 µM, phosHP1α exhibited basal phase separation at low temperature under control conditions, which was also verified with DIC microscopy (Fig. 5). Addition of Mg²⁺ enhanced droplet formation, increasing turbidity at low temperature and extending the stability of the condensed phase to higher temperatures. Across all tested concentrations, turbidity decreased monotonically with increasing temperature, indicating that Mg²⁺-induced phase separation remains favored at low temperature and is progressively destabilized upon heating with a predominantly UCST-like response.

**Figure 5.**
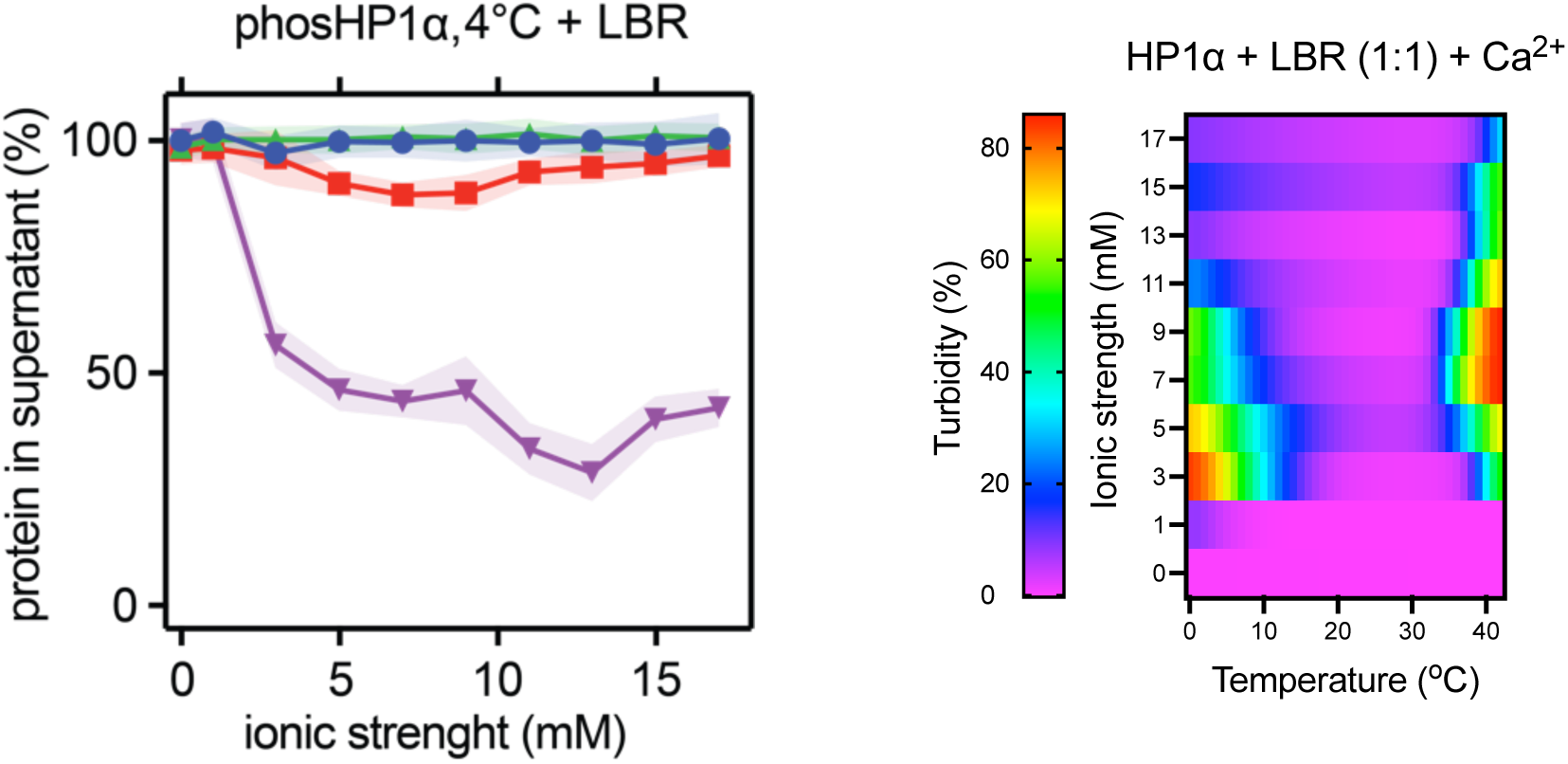
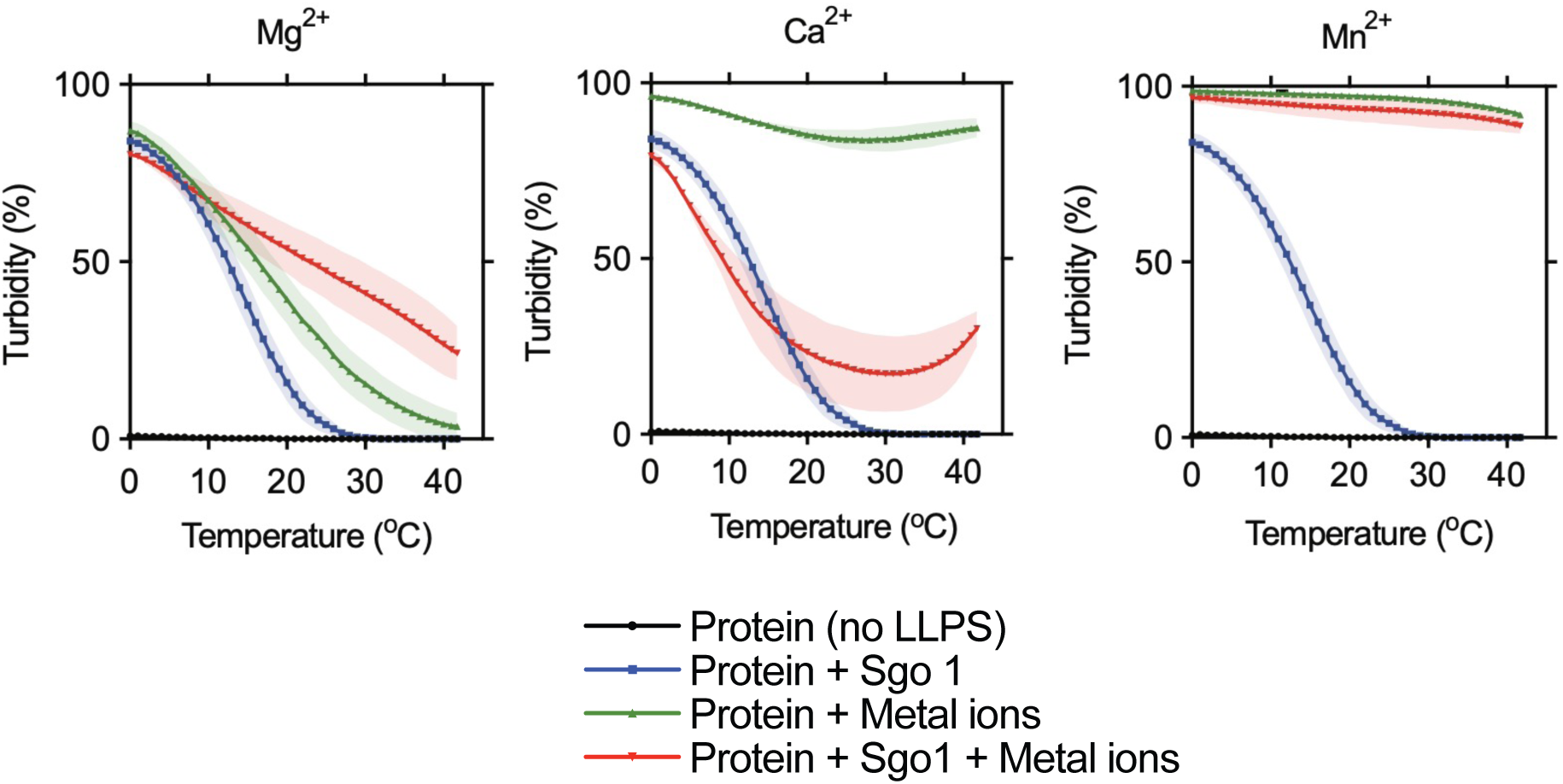
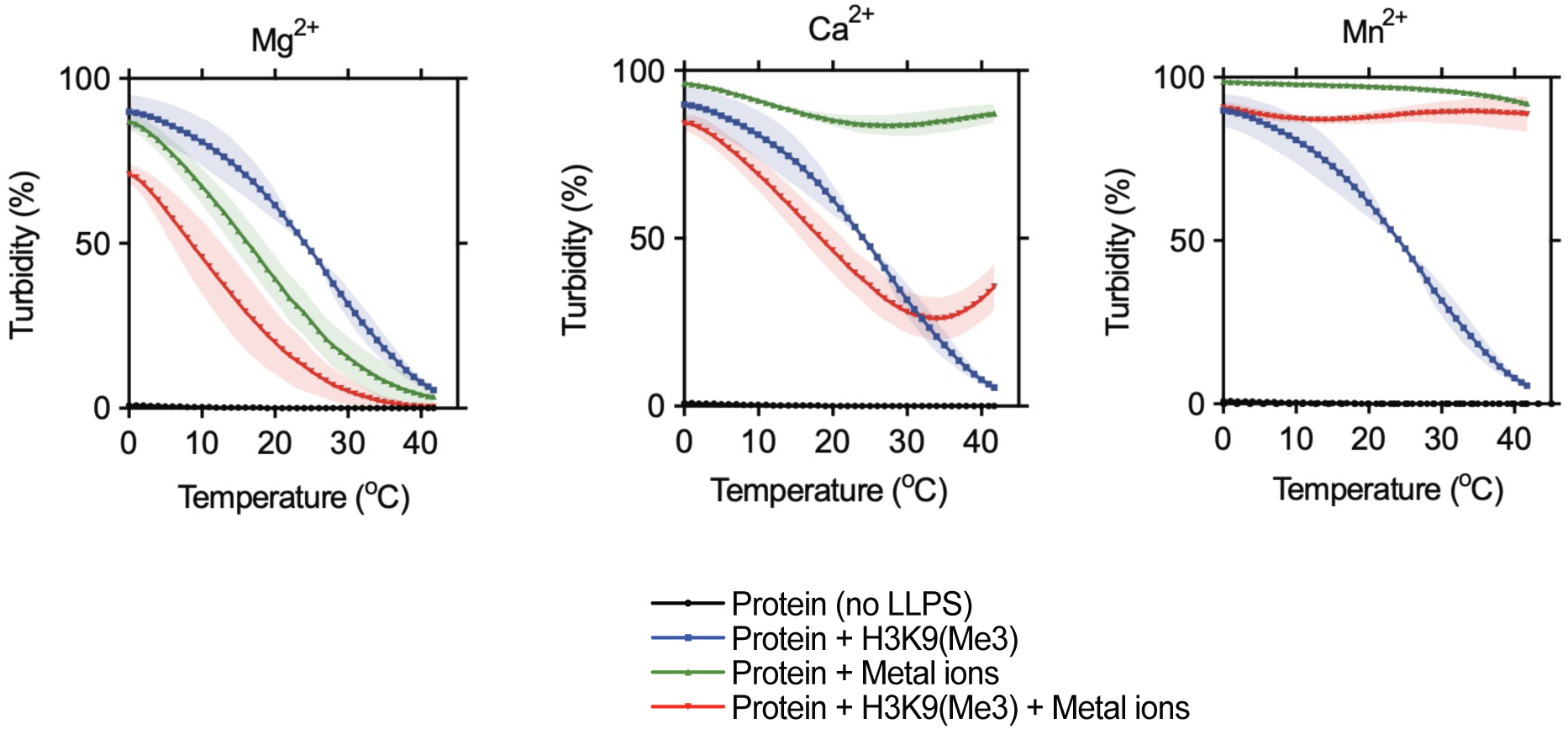
Metal ions modulate the effects of peptide regulators on HP1α phase separation. (A) Effects of the LBR peptide on metal-induced phase separation of phosphorylated HP1α analyzed using depletion assays and turbidity measurements. (B) Temperature-dependent turbidity measurements showing the combined effects of Sgo1-derived peptide and divalent metal ions on HP1α condensate formation. (C) Analogous experiments performed using H3K9me3 peptide. Together, the data demonstrate that metal ions reshape the influence of canonical HP1α interaction partners on condensate formation and stability.

**Figure 6.**
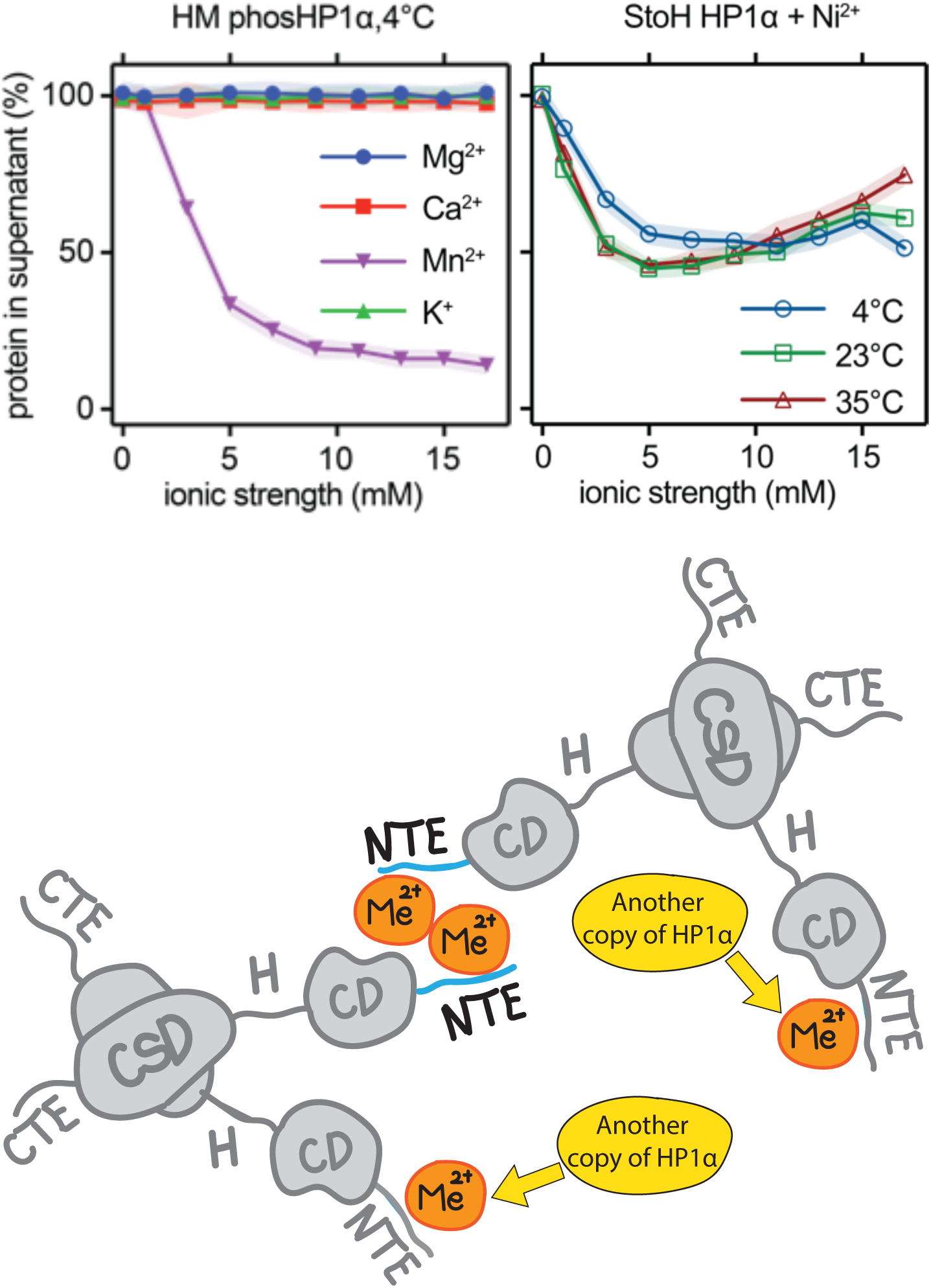
Metal-driven phase separation can occur independently of canonical electrostatic interactions. (A) Phase separation behavior of mutant HP1α variants lacking the native electrostatic interaction network, revealing differential responses to Mg²⁺, Ca²⁺, and Mn²⁺. (B) Engineering of a histidine-rich metal-binding module enables Ni²⁺-dependent phase separation and supports a model in which divalent metal ions promote intermolecular bridging between HP1α molecules. The schematic summarizes the proposed mechanism of metal-mediated intermolecular connectivity within phosphorylated HP1α condensates.

In contrast, Ca²⁺ produced a qualitatively different response. At the lowest concentrations added, turbidity decreased monotonically with temperature, similar to Mg²⁺. At intermediate concentrations (5–9 mM), phase separation was maintained across the entire temperature range, with consistently high turbidity. At higher concentrations, the temperature dependence became non-monotonic, with reduced turbidity at intermediate temperatures and increased phase separation at both low and high temperatures. This behavior indicates that the effect of Ca²⁺ cannot be described by a single thermodynamic regime, but instead reflects the coexistence of distinct interaction modes with opposing temperature dependencies, resulting in features consistent with both UCST– and LCST-like phase behavior.

Mn²⁺ exhibited the strongest effect, inducing robust phase separation even at low concentrations. In contrast to Mg²⁺ and Ca²⁺, manganese supported high turbidity across the entire temperature range, indicating stabilization of the condensed phase under all conditions tested.

Overall, these results demonstrate that divalent metal ions not only promote phase separation of phosHP1α but also reshape its temperature dependence in an ion-specific manner. While Mg²⁺ primarily enhances low-temperature phase separation, Ca²⁺ introduces complex, non-monotonic behavior, and Mn²⁺ strongly stabilizes the condensed phase across all temperatures.

### Metal ions reshape peptide-regulated phase separation

To examine how metal ions influence regulatory interactions, we tested three peptides known to modulate HP1α condensation: the H3K9me3 histone tail peptide, which promotes LLPS through chromodomain binding; the LBR peptide, which inhibits LLPS by binding to the chromoshadow domain; and the Sgo1 peptide, which binds to the same site as LBR but enhances phase separation. Experiments were performed under conditions where phosphorylated HP1α alone showed minimal phase separation, allowing both inhibitory and stimulatory effects to be detected. For LBR and Sgo1, peptide concentrations were chosen to match a 1:1 ratio with the HP1α dimer, whereas H3K9me3 was added at a 1:1 ratio with the monomer, as previously reported (33). Phase behavior was assessed using spin-down assays at 4 °C, and, for selected conditions, temperature-dependent turbidity measurements were performed.

In the absence of metal ions, LBR strongly suppressed phase separation, with all of the protein remaining in the supernatant. Addition of Mg²⁺ partially restored LLPS, reducing the soluble fraction at intermediate concentrations, although this effect remained very limited. In contrast, Ca²⁺ more effectively counteracted LBR inhibition, leading to substantial protein depletion from the supernatant. Temperature ramp experiments further revealed that calcium restores phase separation across a broad temperature range, including conditions where LBR alone fully suppresses droplet formation.

Sgo1, which promotes LLPS, exhibited partially additive behavior with metal ions. In the absence of metals, Sgo1 induced moderate phase separation. Addition of Mg²⁺ or Ca²⁺ further enhanced droplet formation, although the magnitude of this effect was smaller than that observed for LBR rescue. Temperature-dependent measurements indicated that metals primarily extend the stability of Sgo1-induced condensates rather than fundamentally reshaping their behavior.

For the H3K9me3 peptide, which strongly promotes phase separation and mimics the chromatin-bound state of HP1α in a simplified manner, the effect of metal ions was less pronounced. Under conditions where the peptide alone induced robust LLPS, addition of Mg²⁺ or Ca²⁺ led to only modest changes in the fraction of protein partitioning into the condensed phase. However, subtle differences in temperature dependence were observed, suggesting that metal coordination can still modulate condensate stability even when phase separation is already strongly favored.

Interestingly, for Mn²⁺, phase separation of phosHP1α in the presence of the regulatory peptides was dominated by the metal ion, with combined conditions closely resembling those observed for Mn²⁺ alone. As a result, peptide-dependent effects were largely masked under these conditions.

Altogether, these results show that metal ions do not simply enhance or inhibit peptide-driven phase separation, but instead reshape the balance between distinct interaction modes. Depending on the cation, metal-mediated interactions can counteract inhibitory binding, cooperate with droplet-promoting interactions, or compete with canonical binding pathways in a temperature-dependent manner. These findings indicate that metal coordination provides an alternative route for driving condensation that can modulate, and in some cases outweigh, established protein–protein interaction networks.

### Metal-driven phase separation is mechanistically distinct from canonical electrostatic interactions

The results above indicate that divalent metal ions do not simply enhance the canonical phase separation pathway of phosHP1α, but can also promote condensation through an additional mechanism. In the established model, LLPS is driven primarily by electrostatic interactions between the negatively charged phosphorylated NTE and positively charged residues within the hinge. Our data instead suggest that metal coordination can directly contribute to intermolecular association. To test whether these mechanisms can be separated, we designed two complementary perturbations targeting the electrostatic and metal-dependent contributions.

To disrupt the electrostatic pathway, we generated a hinge mutant in which the positively charged KRK motif was replaced with alanines. Under these conditions, phosHP1α no longer undergoes phase separation in the absence of metal ions (30). In spin-down assays at 4 °C, addition of Mg²⁺ or Ca²⁺ failed to restore LLPS, with most of the protein remaining in the supernatant at all tested concentrations (Fig. X). In contrast, Mn²⁺ retained a strong ability to induce phase separation, leading to substantial depletion from the supernatant. These results indicate that Mg²⁺– and Ca²⁺-dependent phase separation relies on the native electrostatic interactions, whereas Mn²⁺ can drive condensation independently of this pathway.

To test whether metal coordination alone is sufficient to induce phase separation, we replaced the phosphoserine cluster with a histidine-based metal-binding motif. In this construct, no LLPS was observed in the absence of metal ions. However, addition of Ni²⁺ induced robust phase separation across all tested temperatures, as reflected by a strong reduction in protein in the supernatant (Fig. X). Unlike the wild-type protein, this metal-induced phase separation displayed little temperature dependence, indicating stabilization of the condensed phase over a broad range of conditions.

These experiments establish that metal-induced phase separation can proceed through a mechanism distinct from the canonical electrostatic pathway. While Mg²⁺ and Ca²⁺ primarily act within the framework of the native phospho-NTE–hinge interaction, Mn²⁺ reveals an alternative mechanism in which metal coordination alone can support intermolecular association. The ability of an engineered histidine-based motif to recapitulate this behavior further demonstrates that multivalent metal binding is, in principle, sufficient to drive condensation. These findings show that phosphorylation can encode not only electrostatic interactions but also latent metal-binding modules, thereby providing an additional tunable layer of regulation for biomolecular phase separation.

## Discussion

Phosphorylation reshapes protein conformational equilibria and molecular interactions by introducing negatively charged phosphate groups. For example, multisite phosphorylation of 4E-BP2 induces folding into a β-structured conformation that enables high-affinity binding to eIF4E (48). In the Sic1 system, multisite phosphorylation converts the disordered protein into a polyelectrostatic ligand, in which multiple weak phosphosites act cooperatively to drive binding (49). A related framework has been proposed for HP1α, where phosphorylation of the N-terminal serine cluster promotes phase separation through interactions between the phosphorylated NTE and basic segments in the chromodomain and hinge region, thereby increasing multivalency and driving oligomerization (30, 50). However, the strong ion-dependent effects observed here indicate that the established electrostatic model does not fully account for HP1α self-association and instead point to an additional interaction pathway contributing to phase separation.

Metal ion interactions are not incidental but are an intrinsic aspect of phosphate chemistry, dictated by its charge, geometry, and multiple oxygen donors. As emphasized by Westheimer, phosphates combine stability in water with persistent ionization, which makes them uniquely suited for biological systems (51). Subsequent analyses show that phosphate chemistry is governed by electrostatics in a broader sense, including interactions with ions and the surrounding environment (52). Consistent with this, phosphate esters readily coordinate mono– and divalent cations, and studies on simple phosphoamino acid analogs have shown that metal binding depends strongly on ion identity, stoichiometry, and ligand geometry (53). In proteins containing closely spaced phosphoserines, these properties enable metal-mediated intermolecular interactions that can organize proteins into higher-order assemblies.

Biological systems exploit the coordination properties of phosphorylated residues to organize macromolecular assemblies with distinct material states. Osteopontin, a highly phosphorylated protein involved in biomineralization, uses its phosphate-rich regions to strengthen interactions with calcium-containing mineral surfaces, modulating crystal growth and contributing to dense, solid-like biomineral structures that are effectively irreversible under physiological conditions (54, 55). Phosvitin, one of the most highly phosphorylated proteins found in nature, illustrates how dense phosphoserine clusters can directly couple metal binding to changes in material state. Multivalent ion coordination reduces electrostatic repulsion between phosphate groups and promotes intermolecular bridging, contributing to compaction and stabilization of dense storage granules in egg yolk (56). In this system, metal coordination stabilizes a kinetically arrested, solid-like, dense phase that is reversible only under sufficiently strong environmental perturbation (57). A particularly instructive example is provided by caseins. These milk proteins contain clusters of closely spaced serine residues that, like human HP1α, are phosphorylated in a hierarchical manner and bind calcium with high capacity through multivalent coordination. Calcium binding drives intermolecular association of caseins into colloidal micellar assemblies that function as a reservoir for calcium and phosphate in milk (21). Depending on ion concentration, temperature, and pH, these assemblies can remain dynamic or transition into more stable, gel-like, or aggregated states (58). Thus, caseins exemplify how clustered phosphoserines can couple to calcium binding to the formation of self-associated states with tunable material properties.

All these systems show that phosphate-mediated metal coordination can drive protein self-association across a continuum of material states, including biominerals, polymeric networks, colloidal micelles, gels, and aggregates. In this context, phosHP1α appears to operate in a distinct regime, in which these interactions remain sufficiently weak and dynamic to support reversible liquid-like condensation. NMR measurements show that interactions between divalent cations and the phosphoserine cluster occur in fast exchange and do not give rise to a well-defined, stoichiometric complex. Instead, the data are consistent with transient coordination events in which ions repeatedly sample the phosphorylated region. This mode of interaction is further supported by the strong PRE effects observed for Mn²⁺ at substoichiometric concentrations, indicating frequent short-lived encounters rather than stable binding. At the macroscopic level, these interactions promote phase separation that is fully reversible upon chelation and highly sensitive to temperature and ion identity. These observations indicate that metal ions act as transient crosslinkers, enhancing intermolecular connectivity without locking the system into a rigid network. As a result, phosHP1α forms liquid-like condensates rather than solid assemblies, placing it in a regime where coordination-driven interactions remain sufficiently labile to support dynamic reorganization of the dense phase.

The extent and nature of this crosslinking depend strongly on the identity of the coordinating metal ion. The ion– and temperature-dependent phase behavior observed here can be rationalized by considering how different cations balance coordination to phosphates with retention or release of their hydration shells. This metal dependence is consistent with earlier studies on simple phosphoamino acid esters, which showed that phosphoserine coordination varies markedly with ion identity, affecting both complex stability and binding geometry (53). Mg²⁺, Ca²⁺, and Mn²⁺ differ not only in ionic radius but also in hydration strength and ligand exchange properties, which together determine whether interactions with phosphoserines occur predominantly through water-mediated (outer-sphere) or more direct (inner-sphere) coordination. Mg²⁺, with its small radius and high charge density, is strongly hydrated and incurs a high energetic penalty for dehydration. Its hydration free energy is substantially larger than that of Ca²⁺ (*ΔG*_hyd_ ≈ −1837 versus −1527 kJ×mol⁻¹), and first-shell water exchange is approximately three orders of magnitude slower, favoring outer-sphere interactions in which water molecules remain interposed between the ion and phosphate groups (59). Such interactions support only weak, transient intermolecular bridging, consistent with the moderate, predominantly UCST-like response observed for Mg²⁺ and phosHP1α. In contrast, Ca²⁺ is larger and more labile, with lower barriers for ligand exchange and a broader range of coordination geometries. As temperature increases and hydration shells become less stable, Ca²⁺ can undergo partial dehydration more readily, enabling direct engagement with phosphate groups (59). This provides a plausible basis for the non-monotonic temperature dependence observed here: at low temperature, the response is dominated by hydration-mediated interactions, whereas at higher temperatures, partial dehydration increasingly promotes intermolecular connectivity, giving rise to coexisting UCST– and LCST-like features.

Unlike alkaline earth metals, Mn²⁺ also possesses partially filled *d*-orbitals, which increase its coordination flexibility and facilitate rapid exchange between distinct phosphate environments. As a first-row transition metal, it offers greater coordination versatility and efficiently samples the phosphoserine cluster, as evidenced by strong PRE effects at substoichiometric concentrations. Once a threshold concentration is exceeded, these interactions likely generate a highly connected network that stabilizes phase separation across the entire temperature range, effectively overriding classical phosHP1α self-association behavior. The behavior of the engineered histidine-rich construct with Ni²⁺ provides an important point of comparison. Because Ni²⁺ strongly favors direct coordination to imidazole donors through ligand-field stabilized interactions, its ability to drive robust condensation demonstrates that multivalent metal coordination is, in itself, sufficient to support dense-phase formation, independently of electrostatic interactions between the phospho-N-terminal region and the hinge. Thus, the UCST– and LCST-like features observed here are not distinct phase separation mechanisms but reflect different temperature sensitivities of hydration-dominated and coordination-driven interactions coexisting within the same system. Overall, the balance between these interaction modes is governed by ion identity, concentration, temperature and competing ligands.

Our results show that phosphorylation expands the interaction landscape of intrinsically disordered proteins by enabling metal coordination in addition to electrostatics. In phosphorylated HP1α, this additional layer supports weak, dynamic intermolecular bridging that drives reversible LLPS and reshapes its thermodynamic behavior in an ion-specific manner. When considered alongside other phosphoproteins, these findings point to a broader mechanism in which phosphate–metal interactions organize proteins into assemblies with distinct material properties. In this framework, the balance between electrostatics, coordination, hydration, and protein dynamics defines a continuum of material states ranging from aggregation to liquid condensation. For chromatin-associated proteins such as HP1α, this mechanism may provide a further route by which local changes in metal ion availability or phosphorylation state contribute to the regulation of higher-order genome organization. This perspective establishes a direct link between post-translational modification, coordination chemistry, and the physical organization of biomolecular matter.

## Methods

### Expression constructs

Human HP1α constructs used in this study were based on the full-length human CBX5 sequence (Uniprot entry P45973). Wild-type codon-optimized synthetic HP1α gene (GenScript) and mutant variants were cloned into bacterial expression vector pET28b containing an N-terminal His-SUMO tag. Mutants used in this study included the KRK→AAA hinge mutant (K90, R91, K92 replaced by alanines), S11A, S14A, S12A/S13A, and the S11–S14→HHHH mutant.

### Protein expression

Proteins were expressed in *Escherichia coli* BL21(DE3) cells transformed with the appropriate expression plasmids. Cells were grown at 37 °C until the optical density at 600 nm reached ∼0.6–0.8, and protein expression was induced with 1 mMIPTG. Expression was continued overnight at reduced temperature (25°C).

For uniformly 13C and 15N-labeled samples used for NMR spectroscopy, cells were grown in D_2_O-based M9 minimal medium supplemented with U-^2^H,^13^C-glucose and ^15^NH_4_Cl as the sole nitrogen source. The M9 medium contained 6 g Na2HPO4, 3 g KH2PO4, 0.5 g NaCl, 1 g NH4Cl, 3 g glucose, 1 mL of 100 mM CaCl2, 1 mL of 1 M MgSO4, thiamine, biotin, kanamycin, and trace element solutions. The pD was adjusted to 7.4 (uncorrected).

### Protein purification

Cells were harvested by centrifugation and resuspended in lysis buffer containing 20 mM phosphate buffer, 500 mM NaCl, 20 mM imidazole, and 5 mM guanidine hydrochloride, pH 7.4. Cells were lysed by sonication and clarified by centrifugation. Proteins were purified using Ni^2+^-affinity chromatography. Washing was performed using a buffer containing 20 mM sodium phosphate, 500 mM NaCl, and 40 mM imidazole, while proteins were eluted using a buffer containing 500 mM imidazole. Following affinity purification, the His-SUMO tag was removed by Ulp1 protease cleavage. Proteins were subsequently dialyzed against 25 mM Tris buffer containing 300 mM NaCl, 1 mM EDTA, and 2 mM DTT, pH 7.4. Final purification was performed by size-exclusion chromatography in 50 mM Tris buffer containing 300 mM NaCl, 2 mM DTT, and 0.01% NaN3, pH 7.2. Final protein samples were concentrated using centrifugal concentrators, flash-frozen in liquid nitrogen, and stored at −80 °C. Protein concentrations were determined spectrophotometrically using calculated extinction coefficients (29450 M^-1^cm^-1^).

### NMR spectroscopy

NMR experiments were recorded at 25°C unless otherwise indicated on spectrometers operating at proton frequencies of 600 (Agilent) and 800 MHz (Bruker). Typical NMR samples contained 100–300 µM protein in 50 mM HEPES buffer, pH 7.0, supplemented with 50 mM KCl and 2 mM TCEP. Samples contained 5–10% D2O for the spectrometer lock. Spectra were processed using NMRPipe and analyzed using Sparky and POKY. Chemical shift perturbations (CSPs) were calculated using standard combined proton/nitrogen weighting schemes:

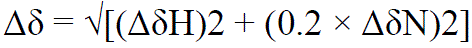

where ΔδH and ΔδN correspond to proton and nitrogen chemical shift differences, respectively.

### HNCO(CA)P experiment pulse sequence scheme

To identify phosphorylated serine residues, a modified HNCoCaP experiment was used. The pulse sequence exploits scalar coupling between the Cα nucleus and the phosphate group of phosphoserine residues. Magnetization transfer proceeds from the amide proton through the peptide backbone to Cα and subsequently to 31P before being transferred back for detection.

The experiment selectively detects residues following phosphorylated serines through the i−1 connectivity pathway. Non-uniform sampling was employed to improve resolution and reduce experimental time.

The HNCA(CA)P experiments were performed according to the pulse sequence scheme. Rectangles represent hard pulses. Filled and empty symbols represent 90° and 180° pulses, respectively. Hard ^1^H, ^13^C, ^15^N and ^31^P π/2 pulses of 10.3, 16.2, 47.4, and 29.6 us were used, respectively. ^1^H, ^2^H, and ^15^N composite pulse decoupling is performed using WALTZ-16 (Shaka et al. 1983) at γB1/2π of 3.7, 0.79, and 1.18 kHz, respectively. Selective CA and CO ^13^C pulses are applied with the RF field strength adjusted to |ΔΩCA-CO|/√15 (√3) for 90° and 180° pulses, respectively. 90° and 180°, rectangular and sinc-shaped pulsed (bell-shaped at the scheme) with a duration of 53.5 (47.9) us and 87.8 (78.4) us, respectively, were used. Off-resonance pulses are applied using phase modulation of the carrier. The PFG durations are set to 0.5 ms, except for coherence selection gradients for which 2.0 ms and 0.2 ms, are used. Delay durations are set as follows: ΔNH=5.4 ms, ΔNCO=28 ms, ΔCOCA=9.1 ms ΔPC=54 ms was optimised to 51 ms. Evolution periods for N and CO are in semi-constant-time mode: ai= (ti+Δ)/2; bi=ti(1– Δ/timax)/2; ci= Δ(1– ti/timax)/2 or in constant-time mode: ai= (ti+Δ)/2; bi=0; ci= (Δ-ti)/2 where Δ stands for ΔNH, ΔNCO. The evolution for 31P in t1 is in real-time mode. The phase cycle is: ϕ_1_=x, –x; ϕ_2_=2x,2(-x), ϕ_3_=4x,4(-x) and ϕ_rec_= ϕ_1_+ϕ_2_+ ϕ_3_ In t1, t2, t3, t4 dimensions quadrature is accomplished using States-TPPI method, by incrementing ϕ_1_, ϕ_2_, ϕ_3_, ϕ_4_ phases, respectively. The phase ψ=x was inverted simultaneously with the last gradient pulse to achieve echo-antiecho coherence transfer selection in the last indirect dimension. 180° water 1.07 ms sinc-shaped flipback pulses are used. All measurements were performed on Agilent DD2 600 spectrometer equipped with 5 mm Penta probes.

### Analysis of metal titration data

Because the phosphoserine region constitutes a multivalent interaction module with potentially heterogeneous binding modes, the titration data were analyzed using empirical saturating functions rather than simple 1:1 binding models. Apparent EC50 values were obtained from global fitting procedures.

### Paramagnetic relaxation enhancement experiments

Paramagnetic relaxation enhancement (PRE) experiments were performed using MnCl_2_. Small amounts of Mn^2+^ were added to phosphorylated and non-phosphorylated HP1α samples, and resonance intensity attenuation was monitored in ^15^N HSQC spectra.

### Turbidity measurements

LLPS experiments were performed using phosphorylated HP1α diluted into reference buffer containing 20 mM HEPES, 75 mM KCl, and 5 mM DTT, pH 7.4. Turbidity measurements were carried out using a Cary 3500 Agilent UV–Vis spectrophotometer. For experiments with Mg^2+^, 200 µM phosHP1α was used, whereas experiments with Ca^2+^ and Mn^2+^ employed 100 µM phosHP1α. Protein samples were incubated for 10 min at 37 °C in PCR tubes before transfer to quartz cuvettes. Metal ion stock solutions were then added directly to the cuvette. Samples were incubated for 10 min at 0 °C before initiating temperature ramps. The temperature was increased at a rate of 1 °C/min until reaching 42 °C, with turbidity measurements collected every minute. Turbidity was monitored at 600 nm. Control samples contained buffer without added metal ions. Each experiment was repeated at least three times. To facilitate comparison between mono– and divalent salts, ionic strength contributions were calculated according to:

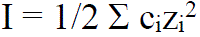

where c_i_ corresponds to ion concentration and z_i_ to ion charge.

### Spin-down assays

Phase separation was quantified using centrifugation-based depletion assays. Protein samples were incubated under LLPS conditions at the indicated temperatures (30 minutes) and centrifuged (10 minutes at the same temperature, 10 000 g) to separate dilute and dense phases. Following centrifugation, the concentration of protein remaining in the supernatant was determined spectrophotometrically by absorbance at 280 nm. Protein depletion was expressed relative to control samples lacking metal ions. Ionic strength contributions of salts were calculated according to I = 1/2 Σc_i_z_i_^2^, where c_i_ corresponds to ion concentration and z_i_ to ion charge.

### Peptide modulation experiments

Synthetic peptides corresponding to HP1α interaction partners, including H3K9me3-derived peptides, Sgo1-derived peptides, and LBR-derived peptides, were added to phosphorylated HP1α samples prior to LLPS measurements. The effects of peptides on phase separation were analyzed in the absence and presence of divalent metal ions using turbidity assays and spin-down experiments as described above.

